# Inhibition and late errors in the antisaccade task: Influence of task design

**DOI:** 10.1101/270165

**Authors:** Eduardo A. Aponte, Dominic G. Tschan, Klaas E. Stephan, Jakob Heinzle

## Abstract

In the antisaccade task, subjects are instructed to saccade in the opposite direction of a peripheral visual cue (PVC). Importantly, several psychiatric disorders are associated with increased error rates in this paradigm. Despite this observation, there is no consensus about the mechanism behind antisaccade errors: while often explained as inhibition failures, some studies have suggested that errors are caused by deficits in the ability to initiate voluntary saccades. Using a computational model, we recently showed that under some conditions high latency or late errors can be explained by a race process between voluntary pro- and antisaccades. A limitation of our findings is that in our previous experiment the PVC signaled the trial type, whereas in most studies, subjects are informed about the trial type before the PVC is presented. We refer to these task designs as *asynchronous* (*AC*) and *synchronous cues* (*SC*) conditions. Here, we investigated to which extent differences in design affect the type and frequency of errors in the antisaccade task. Twenty-four subjects participated in mixed blocks of pro- and antisaccade trials in both conditions. Our results demonstrate that error rates were highly correlated across task designs and a non-negligible fraction of them were classified as late errors in both conditions. In summary, our findings indicate that errors in the AC task are the result of both inhibition failures and inaccurate voluntary action initiation.

## Introduction

The antisaccade task (Hallett, 1978) is an oculomotor paradigm widely used in psychiatric research (Hutton and Ettinger, 2006; Gooding and Basso, 2008; Bittencourt et al., 2013), in which participants are required to saccade in the opposite direction of a peripheral visual cue. This paradigm probes both the ability to inhibit reflexive responses - i.e. (pro)saccades towards a visual cue- and to initiate voluntary actions -i.e. (anti)saccades in the contralateral direction of the peripheral stimulus (Everling and Fischer, 1998). The clinical relevance of this paradigm originates from the fact that error rates (ER) and reaction times (RT) are altered in many psychiatric and neurological diseases. For example, ERs are higher not only in schizophrenic patients (Gooding and Basso, 2008), but also in their first order relatives as well as in related psychiatric populations, such as in schizoaffective disorder patients (Calkins et al., 2004; Reilly et al., 2014; Myles et al., 2017).

Error in this task have often been attributed to deficits in inhibitory control (e.g. Levy et al., 1998; Broerse et al., 2001; Calkins et al., 2004), but these have been also related to deficits in voluntary action initiation. Initially, Fischer et al., 2000 proposed to differentiate between inhibition and volitional errors by highlighting the large discrepancies in performance between gap and overlap paradigms. Interestingly, in a large cohort, Fischer et al., 2000 showed that subjects that mainly perform ‘express’ errors tend to correct their mistakes more often than participants that produce mostly late errors. Using a similar argument, Klein and Fischer, 2005 proposed to extend the distinction between express and ‘normal-range’ saccades to antisaccade errors, and used indirect statistical evidence to suggest that these evolve differently during development and are correlated with different psychometric constructs (Klein et al., 2010). Reuter and colleagues (Reuter and Kathmann, 2004; Reuter et al., 2005), based on the parallel programming model proposed by Massen, 2004, hypothesized that at least some fraction of the errors observed in this paradigm are caused by failures to initiate a voluntary action. More recently, Lo and Wang, 2016 incorporated the idea of two types of errors into a biophysical model of eye movement control and speculated that the mechanisms behind late errors might be of interest in psychiatric research. In that spirit, Coe and Munoz, 2017 suggested that the ratio between early and late errors could distinguish between control and patient populations, such as Parkinson’s disease and lateral amyotrophic sclerosis patients.

Recently (Aponte et al., 2017), using the *Stochastic Early Reaction, Inhibition, and late Action* (SERIA) model, we presented quantitative and qualitative evidence that errors in the antisaccade task can be divided in fast, reflex-like prosaccades and voluntary but erroneous late prosaccades. SERIA is a formalized extension of the model initially proposed by Noorani and Carpenter, 2013 (but see Camalier et al., 2007; reviewed in Cutsuridis, 2017; Noorani, 2017) and builds on the idea that RTs are distributed as the threshold hit times of linear, ballistic accumulation processes (Noorani and Carpenter, 2016). In this family of models, pro- and antisaccades are generated by two competing but independent accumulators. In addition, a third unobservable process can stop reflexive prosaccades, similarly as in the model used of the counter demanding saccade task (Logan et al., 1984).

Conceptually, SERIA extends Noorani and Carpenter’s work by introducing a further decision process that can generate late prosaccades and competes with the antisaccade process. Errors can be therefore divided into early errors, explained as inhibition failures, and late errors, explained as the result of a late race between voluntary pro- and antisaccades. Moreover, according to SERIA, errors in prosaccade trials occur when an early response is inhibited, but an antisaccade overwrites a late prosaccade. Thus, our model provides a unified account of all types of errors in the antisaccade task.

One limitation of the study reported in Aponte et al., 2017 is that the version of the antisaccade task used there has not been extensively investigated in humans (in the macaque monkey see e.g. Sato and Schall, 2003; in humans see Weber, 1995; Irving et al., 2009; Liu et al., 2010; Chiau et al., 2011; Weiler and Heath, 2012). Concretely, in Aponte et al., 2017 subjects performed interleaved pro- and antisaccade trials, in which a peripheral stimulus signaled the trial type and the target location (see Fig 1A). We refer to this version of the antisaccade task as a *synchronous cue* (SC) design.

In humans, the antisaccade task is most often administered in a block design (Antoniades et al., 2013) in which subjects perform a single trial type throughout a block and are informed about the task in advance. Even when different trial types are interleaved, participants are usually informed about the task demands before the peripheral cue is presented (e.g., Cherkasova et al., 2002; Massen, 2004; O’Driscoll et al., 2005; Reuter et al., 2006; Pierce et al., 2015; Pierce and McDowell, 2016a; 2016b). For examples of this task in the monkey literature see e.g. (Amador et al., 1998; Johnston et al., 2014; Koval et al., 2014; Vijayraghavan et al., 2016). We refer to this paradigm as the *asynchronous cues* (AC) design.

The main goal of the present study was to test whether the conclusions drawn in our previous study hold for the most commonly used version of the antisaccade task. We acquired data from twenty-four participants in both the SC and AC conditions and compared RT and ER as well as model parameters estimated from the data. We were interested in three main questions: First, we investigated whether in an AC design it was necessary to postulate a late race between voluntary pro- and antisaccades. Hence, we compared models that incorporated a late race against models in which all late saccades were antisaccades. Second, we were interested in differences in the probability of inhibition failures and late errors in different task designs. In particular, we investigated if and in what proportions late errors occurred in SC and AC tasks. Finally, we tested whether the effects of trial type probability reported in Aponte et al., 2017 could be replicated, and whether these effects generalize to the AC design.

## Methods

### Participants

Twenty-five healthy male volunteers (age: 21.4±2.0 y) participated in the study approved by the local ethics board of the Canton of Zurich, Switzerland (KEK-ZH-Nr.2014-0246) and conducted according to the Declaration of Helsinki. Because this experiment was part of a larger pharmacological study, only male participants were included. All subjects had normal or corrected to normal vision and gave their written informed consent to participate.

### Apparatus

The experiment took place in a dimly illuminated room. Subjects viewed a CRT screen (41.4×30cm; Philips 20B40) operating at 85Hz from a distance of 60cm, while their gaze was recorded using an Eyelink 1000 (SR Research, Ottawa, Canada). Head position was stabilized using a chin rest. Gaze position was stored at a sampling rate of 1000Hz. Every block started with a 5-points calibration procedure. Absolute calibration error was aimed to be below 1°. The experiment was programmed in the Python programming language (2.7) using the *PsychoPy* (1.82.02) package (Peirce, 2007; 2008). The experiment was controlled by a personal computer (Intel Core i7 4740K) equipped with a Nvidia GTX760 graphics card.

### Experimental design

The experimental design used here is an extension of the design used in Aponte et al., 2017. Subjects participated in 6 blocks of mixed pro- and antisaccade trials. Each block consisted of 200 randomly interleaved pro- and antisaccade trials, from which either 20, 50 or 80% were prosaccade trials. In addition to trial type probability, we also manipulated the order in which the trial type cue and the saccade direction cue were presented: Subjects were either simultaneously informed about the trial type and saccade direction using one peripheral cue (SC condition), or they were signaled about the trial type before being presented with the peripheral cue (AC condition). Both conditions are explained in detail below. All instructions were given to the participants in written format.

The experiment followed a with-in subject, 3×2 factorial design, with factors *prosaccade trial probability* (PP) with levels PP20, PP50, and PP80 and *cue type* (CUE) with levels SC and AC. The blocks belonging to one of the CUE conditions were administered consecutively. The order of presentation of the blocks was pseudo-randomized and counterbalanced across subjects. The same sequence of pro- and antisaccade trials was used for each PP condition independently of the CUE condition. The peripheral cue was presented randomly on the right and left side of the screen. Again, the same random sequence was used across subjects.

Before participating in the main experiment, subjects underwent a training block for each condition. These consisted of 100 trials, from which the first half were prosaccade trials, followed by 50 antisaccade trials. During training, participants received automatic feedback after each trial indicating whether they had saccade in the correct direction. In order to urge participants to respond speedly, saccades with a latency above 500ms were signaled as errors.

### Synchronous cue (SC) condition

Throughout the experiment, two red circles of 0.25° of radius were presented at ±12°. Each trial started with a cross (0.6×0.6°) displayed at the center of the screen. Subjects were required to fixate for at least 500ms. If their gaze drifted outside a 3° window, the fixation interval was restarted. The fixation target was presented for a further random interval (500-1000ms), after which a green bar (3.48×0.8°) centered on one of the peripheral red circles was displayed for 500ms (Fig. 1A). The bar was presented in either horizontal or vertical orientation. Subjects were instructed to saccade towards the red circle ipsilateral of a horizontal bar, or to the contralateral circle in the case of a vertical bar. The next trial started 1000ms after the peripheral cue was removed.

### Asynchronous cues (AC) condition

The start of the AC condition (Fig. 1B) was identical as in the SC condition, but after the initial fixation period a green bar (3.48×0.8°) was displayed for 700ms, centered on the fixation cross. The bar could be in horizontal or vertical orientation, cueing a pro- or antisaccade trial, respectively. The fixation cross and the green bar were removed at the end of the 700ms period and subsequently a green square (1.74×1.74°) was presented on one of the peripheral red circles for 500ms. Subjects were instructed to saccade to the circle ipsilateral to the green square if the task cue was a horizontal bar, and to saccade to the contralateral circle if it was a vertical bar. The next trial started 1000ms after the green square was removed.

**Fig. 1:**
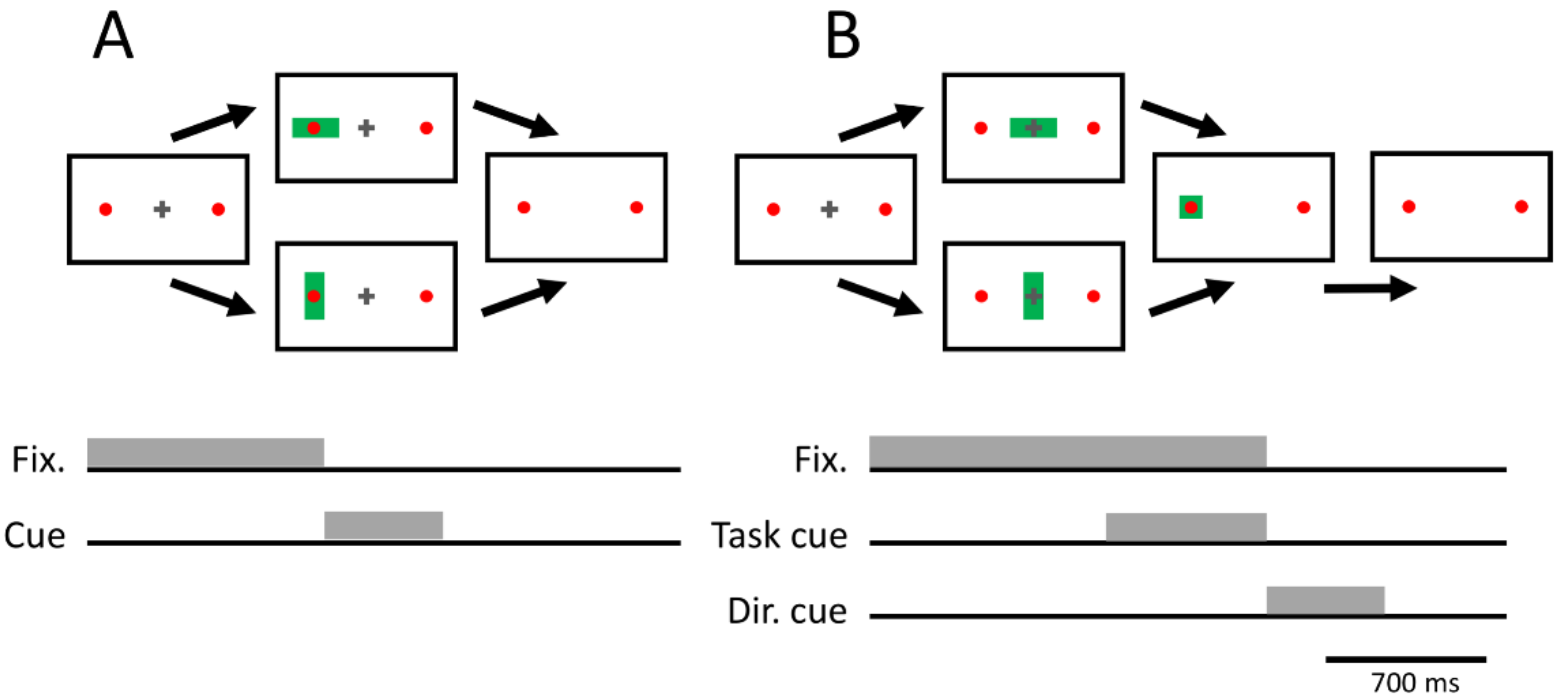
Task design. **A.** *Synchronous cue (SC) condition.* Similarly to Aponte et al., 2017, subjects were instructed to fixate a central cross for 500-1000ms, while two red circles (0.25° radius) were displayed at ±12°. Immediately after the fixation period, a green bar (3.4×0.8°) was displayed centered on one of the red circles for 500ms. Participants were instructed to saccade as fast as possible to the red circle ipsilateral to a horizontal green bar, and to saccade to the contralateral circle when a vertical bar was displayed. **B.** *Asynchronous cues (AC) condition*. As in the SC condition, subjects were instructed to fixate a central cross for 500 to 1000ms. After the initial fixation period, a green bar (3.4×0.8°) was displayed at the center of the screen for 700ms. Immediately afterwards, the fixation cross and the green bar were removed and a green square (1.74×1.74°) was displayed centered on one of the circles. Subjects were instructed to saccade to the circle ipsilateral to the peripheral cue if a horizontal bar was presented, and to saccade to the contralateral circle otherwise.

### Data preprocessing

Data was preprocessed using the Python programming language (2.7). Saccades were detected using the algorithm provided by the eye tracker manufacturer (Stampe, 1993), which uses a velocity and acceleration thresholds of 22°/s and 3800°/s^2^, respectively. Saccades with a magnitude lower than 2° were ignored. RT was defined as the latency of the first saccade after the fixation cross was removed (henceforth, the *main saccade*). Trials were discarded if any of the following conditions was true: if a blink occurred between the start of the fixation period and the end of the main saccade; if subjects failed to maintain fixation; if a saccade had a latency above 800ms or below 50ms, and in the case of an antisaccade, a latency below 95ms. Corrective antisaccades were defined as saccades contralateral to the peripheral cue that followed an error in an antisaccade trial. Corrective saccades were only included in the analysis if they occurred within 900ms after cue presentation and if their end location was within a 4° window around the correct target.

### Classical statistical analysis

Mean RTs, ERs and parameter estimates of the model (see below) were analyzed using a generalized mixed effects linear model. The independent variables were PP with levels PP20, PP50, PP80; CUE with levels SC and AC; SUBJECT entered as a random effect, and, when pro- and antisaccade trials were analyzed together, *trial type (TT*). All regressors were entered as categorical variables. ERs were analyzed using a binomial regression model with the probit function as link function. When probabilities were analyzed, a fixed effects Beta regression model (Cribari-Neto and Zeileis, 2009) was used, because a mixed effect model proved numerically unstable. For RT, we report tests based on the F-statistic, whereas for ER and probabilities we report tests based on the *X*^2^ statistic. Statistical significance was asserted at α=0.05. All statistical tests were performed with the *R* programming language (3.4.2) using the functions *lmer*, *glmer*, and *glmmadmb* (Beta regression model) from the packages *lme4*, *lmerTest*, and *glmmADMB*.

### Modeling

Two models (described in detail in Aponte et al., 2017) were fitted to actions (pro- or antisaccades) and RTs. First, we fitted the *PRO-*, *Stop and Antisaccade* (PROSA) model, which structurally resembles the model described in Noorani and Carpenter, 2013. According to it, three linear race decision units determine RTs and ERs in the antisaccade task. Each unit triggers or stops different types of action depending on the order and time at which they hit threshold (henceforth *hit time*): The first *early unit* triggers a prosaccade if it hits threshold before all other units. These fast reactions can be stopped by the inhibitory unit, if the latter hits threshold before the *early unit*, in analogy to the model originally proposed by Logan et al., 1984. If an early response is inhibited, the third unit triggers an *antisaccade* once it hits threshold. This model represents the hypothesis that all voluntary or late responses are antisaccades.

Second, we fitted the SERIA model, which extends the PROSA model by including a fourth unit that can trigger late, voluntary prosaccades. Hence, SERIA distinguishes between reflexive, early prosaccades, and voluntary late prosaccades. This leads to two types of antisaccade errors: inhibition failures, when the early unit hits threshold before all other units, and volitional or late errors when the late prosaccade unit hits threshold before the antisaccade unit. An error in a prosaccade trial occurs when an early response is stopped, but the antisaccade unit hits threshold before the late prosaccade unit. Note that the model used here corresponds to the SERIA model with late race (SERIA_lr_) introduced in (Aponte et al., 2017).

To fit the models to empirical data, we evaluated three different parametric distributions for the increase rate (or reciprocal hit time) of each of the units: Either we assumed that the increase rate of all the units were truncated Gaussian distributed in analogy to the LATER model (Noorani and Carpenter, 2016), or that the increase rates of the early and inhibitory unit were Gamma distributed, but the increase rate of the late units was inverse Gamma distributed. We refer to this model as the mixed Gamma model. Finally, we considered a model in which the increase rate of all the units was Gamma distributed. In addition, we assumed that a different set of parameters for each of the units was necessary for each trial type. However, we also considered a constrained version of the SERIA model in which the early and inhibitory units followed the same distribution in pro- and antisaccade trials, but the late units had different parameters across trial types (Aponte et al., 2017). For PROSA, we investigated a model in which the early unit followed the same distribution across trial types but others were allowed to differ(Noorani and Carpenter, 2013; Aponte et al., 2017). A summary of the model space is presented in Table 1. Details regarding the number of parameters can be found in Aponte et al., 2017.

**Table 1:** Model space. **List of models with corresponding increase rate distributions and number of free parameters**. In constrained models, some of the parameters are assumed to be equal across trial types. Note that besides the parameters of the units, all models include three nuisance parameters that account for no-response time, late response cost, and the frequency of outlier, i.e., saccades with latencies below the no-response time. Further details can be found in Aponte et al., 2017.

| Model | Parametric dist. | No. parameters |
| --- | --- | --- |
| Unconstrained/Constrained |  | **PROSA** |
| m<sub>1</sub>/m<sub>2</sub> | Truncated Normal | 15/13 |
| m<sub>3</sub>/m<sub>4</sub> | Mixed Gamma | 15/13 |
| m<sub>5</sub>/m<sub>6</sub> | Gamma | 15/13 |
|  |  | **SERIA** |
| m<sub>7</sub>/m<sub>8</sub> | Truncated Normal | 19/15 |
| m<sub>9</sub>/m<sub>10</sub> | Mixed Gamma | 19/15 |
| m<sub>11</sub>/m<sub>12</sub> | Gamma | 19/15 |

We fitted the data of all subjects and PP conditions simultaneously using a Bayesian hierarchical model (Gelman et al., 2003), in which the prior distribution of the parameters for each subject was informed by the population distribution. However, the two CUE conditions were fitted independently, because our goal was to evaluate whether different models were favored under different task designs. The population distribution was modeled using a linear mixed effects model with PP as fixed effect and SUBJECT as a random effect. Details are provided in the Supplementary Methods and Supplementary Figure S1.

Models were fitted using the Metropolis-Hastings algorithm. Their evidence was computed with thermodynamic integration (Gelman and Meng, 1998, Aponte et al., 2016), with 32 chains and a 5^th^ order temperature schedule (Ben Calderhead and Girolami, 2009). To increase the efficiency of the algorithm, we incorporated a ‘swap-step’ according to population MCMC’s accept/reject rule (Ben Calderhead and Girolami, 2009). The algorithm was run for 16 × 10^4^ iterations, but the first 6 × 10^4^ samples were discarded as ‘burn-in’ samples. The code was executed on a computer cluster running Linux (CentOS 7.4.1708), MATLAB R2015a (8.5.0.197613), and GSL 1.16. The software implemented here is publicly available as part of the TAPAS toolbox (see software note). The statistic used to compare models was the difference in log model evidence (LME), which correspond to log Bayes factors (Kass and Raftery, 1995). Because our main hypothesis was related to families of models (SERIA and PROSA), we used Bayesian family model comparison (Penny et al., 2010) implemented in the SPM12 software package (release 6470, function *spm_compare_families.m*). This method pools the evidence of models which are assumed to belong to the same family and returns the posterior probability of each family.

## Results

Twenty-four subjects were included in the final analysis. One subject was excluded because of incomplete data. A total of 28815 main saccades were collected, from which 1079 or 3.7% were discarded. Regarding corrective saccade, 983 and 696 trials were included in the analysis of the SC and AC conditions, respectively.

### Error rate (ER)

Fig. 2A and 2B display the mean ER in all conditions and trial types. Pro- and antisaccade ERs were submitted to two independent tests using PP and CUE as explanatory variables. ERs were higher in the SC condition, regardless of trial type (pro. trials: X^2^(2, *N* = 144) = 402.75, *p* < 10^−5^, anti. trials: X^2^(2, *N* = 144) = 257.06, *p* < 10^−5^). Moreover, there was a significant interaction between the PP and CUE factors in both trial types, demonstrating that PP had a much more pronounced effect in the SC condition (pro. trials: X^2^(2, *N* = 144) = 43.00, *p* < 10^−5^; anti. trials: X^2^(2, *N* = 144) = 63.43, *p* < 10^−5^). Next, we submitted ERs in the two CUE conditions to two separate tests with explanatory variables TT and PP. Thus, we could test whether PP had a significantly different effect in pro- and antisaccade trials. We found that in the two CUE conditions, the interaction between PP and TT was significant (SC: X^2^(2, *N* = 144) = 700.46, *p* < 10^−5^, AC: X^2^(2, *N* = 144) = 41.24, *p* < 10^−5^).

**Fig. 2:**
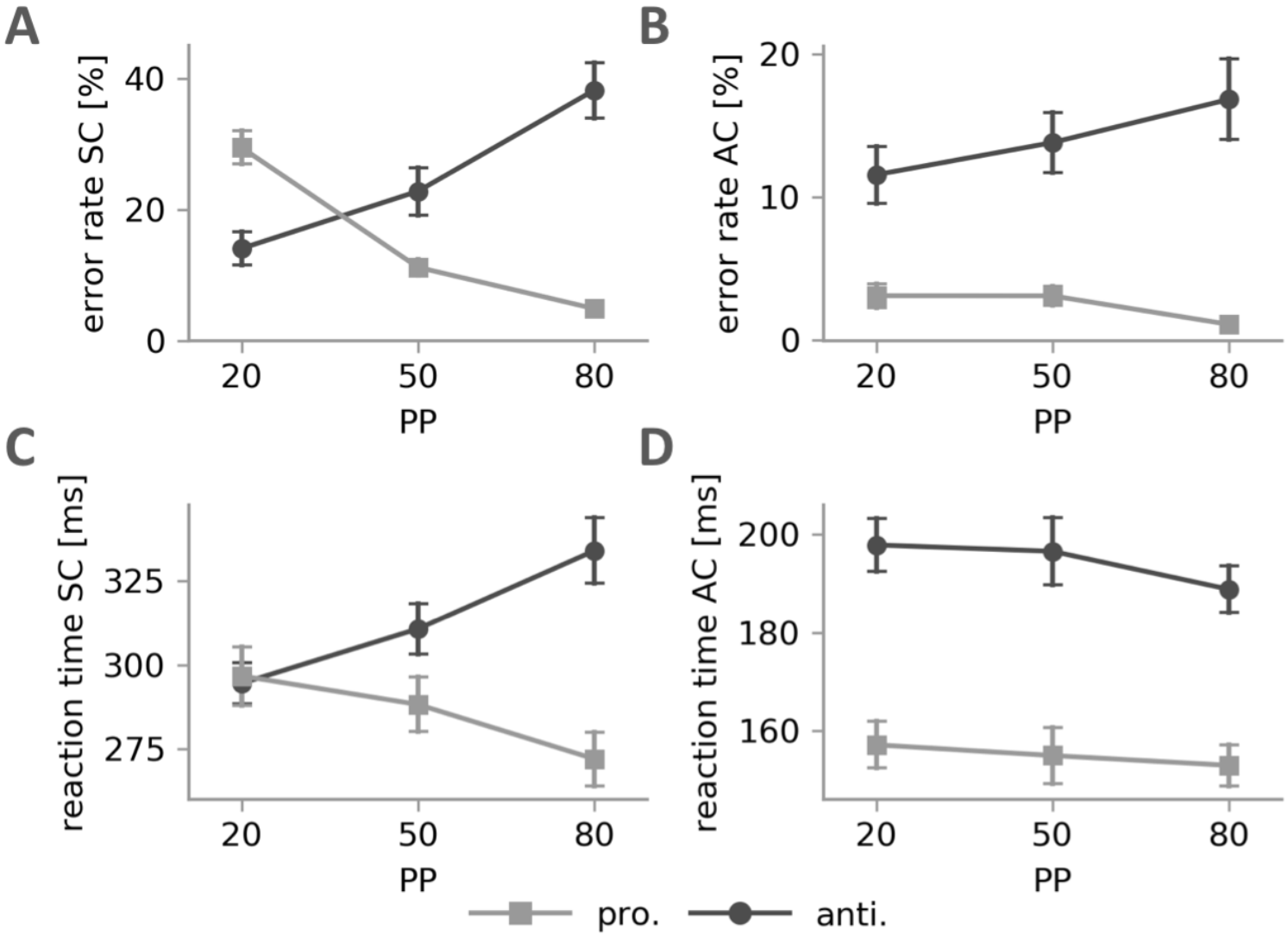
**A.** Mean ER, SC condition. **B.** Mean ER, AC condition. **C.** Mean RT, AC condition. **D.** Mean RT, AC condition. Only the mean RTs of correct trials are displayed. Error bars depict the sem.

Next, we investigated the intra-subject correlation between ER across the two CUE conditions (Fig. 2C). The probit transformed ERs in each PP block were analyzed separately. For numerical reasons, zero percent ERs were inflated to the ER corresponding to a single error. There was a significant correlation between ERs in antisaccade trials for all three PPs (*p* < 0.001), but we found no comparable results in prosaccade trials.

**Figure 3:**
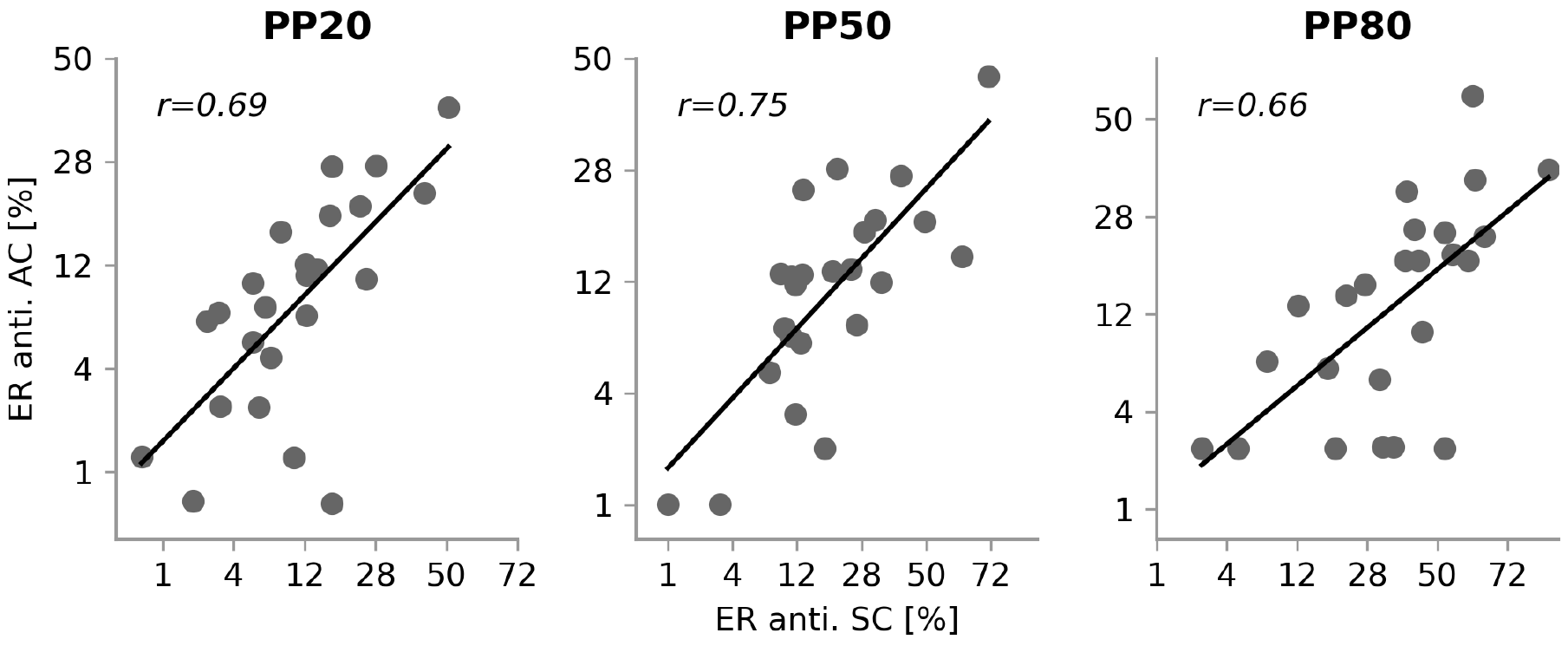
Error rate (ER) in antisaccade trials in the SC and AC conditions. ER were probit transformed.

### Reaction times (RT)

Mean RTs of correct saccades are displayed in Fig. 2B. First, RT in pro- and antisaccade trials were submitted to two separate models with PP and CUE as independent variables. Clearly, RTs in the SC condition were much higher than in the AC condition (pro.: *F*_1,115_ = 815.05, *p* < 10^−5^; anti.: *F*_1,115_ = 789.90, *p* < 10^−5^). The factor PP was significant in both pro- (*F*_2,115_ = 3.46, *p* = 0.03) and antisaccade trials (*F*_2,115_ = 4.32, *p* = 0.01). However, there was a significant interaction between the factors CUE and PP in antisaccade (*F*_2,115_ = 11.25, *p* < 10^−3^), but not in prosaccade trials (*F*_2,115_ = 1.79, *p* = 0.17).

We then investigated both CUE conditions separately in a model with factors PP and TT. In the AC condition, both pro- and antisaccade RTs decreased with PP, as previously reported by (Pierce et al., 2015). However, neither the main effect of PP (*F*_1,115_ = 2.40, *p* = 0.09) nor the interaction PP*TT was significant (*F*_2,115_ = 0.48, *p* = 0.61), although but the main effect of TT was significant (*F*_1,115_ = 238.93, *p* < 10^−5^). In the SC condition, PP had the opposite effect on pro- and antisaccades which resulted in a significant interaction between PP*TT (*F*_2,115_ = 12.99, *p* = 10^−5^).

### Model comparison

In order to compare models, we used the differences in (LME) or log Bayes factors between the hierarchical models fitted to our data (Table 2). The expected log likelihood or accuracy of each model is reported in Supplementary Table S1. This measure is closely related to the *R*^2^ statistic and reflects the un-penalized goodness of fit of a model.

**Table 2:**
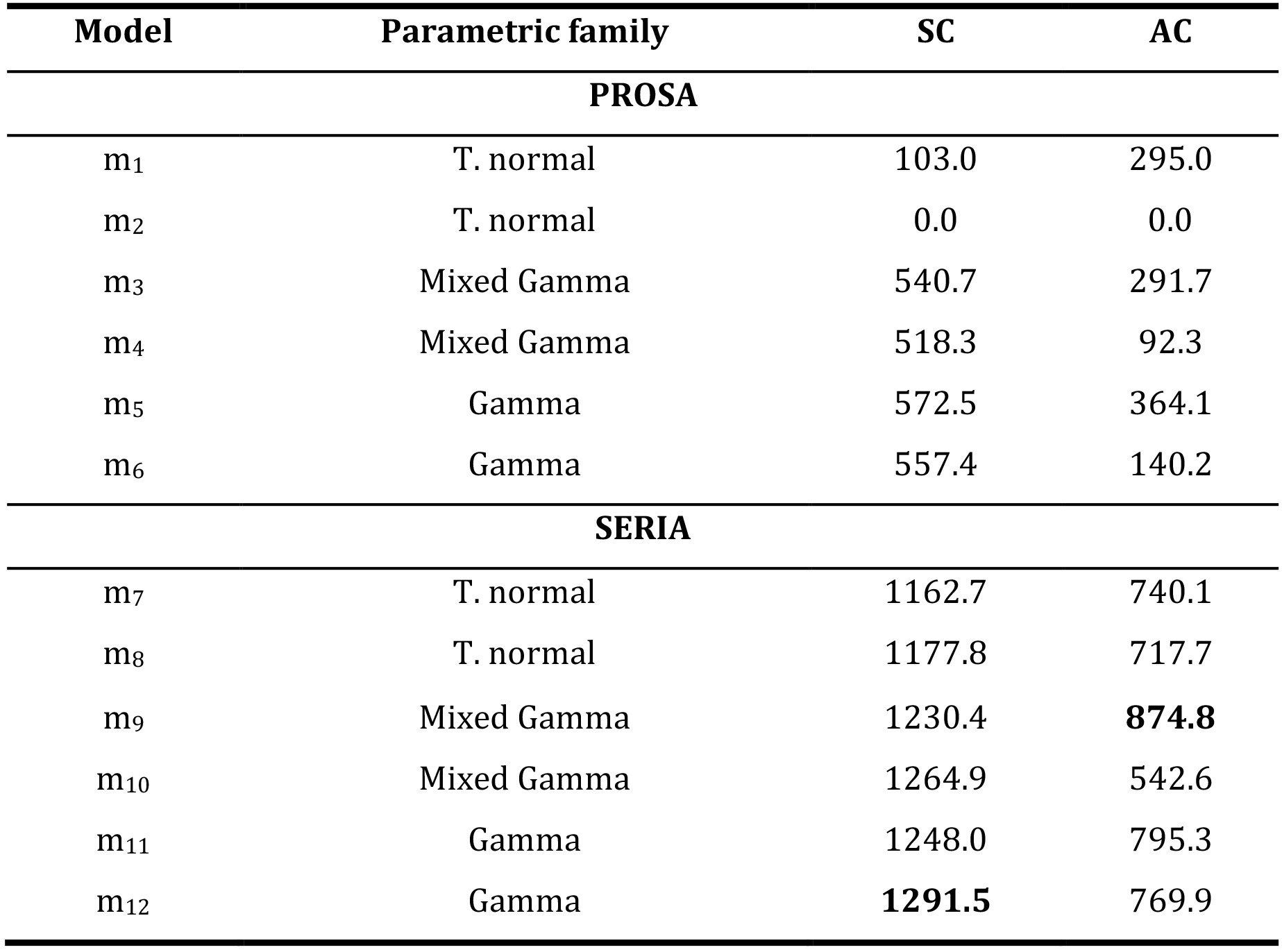
Differences in log model evidence (LME) **Model comparison**. Log evidences were baselined using the lowest value in each condition. The models with the highest evidence are high lightened in bold font.

| Model | Parametric family | SC | AC |
| --- | --- | --- | --- |
| **PROSA** |  |  |  |
| m<sub>1</sub> | T. normal | 103.0 | 295.0 |
| m<sub>2</sub> | T. normal | 0.0 | 0.0 |
| m<sub>3</sub> | Mixed Gamma | 540.7 | 291.7 |
| m<sub>4</sub> | Mixed Gamma | 518.3 | 92.3 |
| m<sub>5</sub> | Gamma | 572.5 | 364.1 |
| m<sub>6</sub> | Gamma | 557.4 | 140.2 |
| **SERIA** |  |  |  |
| m<sub>7</sub> | T. normal | 1162.7 | 740.1 |
| m<sub>8</sub> | T. normal | 1177.8 | 717.7 |
| m<sub>9</sub> | Mixed Gamma | 1230.4 | **874.8** |
| m<sub>10</sub> | Mixed Gamma | 1264.9 | 542.6 |
| m<sub>11</sub> | Gamma | 1248.0 | 795.3 |
| m<sub>12</sub> | Gamma | **1291.5** | 769.9 |

We first compared families of models in each of the conditions separately. In the SC condition, the SERIA family was favored when compared to the PROSA family (posterior probability approx. 1). In the SERIA family, constrained models were favored when compared to models in which the early and inhibitory unit were allowed to differ across trial types (posterior probability approx. 1). When we considered each model independently, analogously to the findings in Aponte et al., 2017, a constrained SERIA model (m_12_) obtained the highest evidence (*ΔLME* > 26.6).

In the AC condition, while the SERIA model was favored when compared to the PROSA model (posterior probability approx. 1), SERIA models in which the early and stop units were not constrained obtained the highest evidence (posterior probability approx. 1). The unconstrained mixed Gamma SERIA model (m_9_) was favored among all possibilities (*ΔLME* > 79.5).

In order to facilitate the comparison across the CUE conditions and our previous study (Aponte et al., 2017), in the following we report the parameter estimates obtained using mixed Gamma models (SC condition:m_10_; AC condition:m_9_).

### Model fits

To qualitatively evaluate the PROSA and SERIA models (Gelman et al., 2003; Gelman and Shalizi, 2013), in Fig. 4 we plotted the histogram of RTs of all saccades and the fit of the best model in each family. For the PROSA model we used model m_5_ in both conditions. Fits were computed by weighting the expected probability density function. in a given block by the corresponding number of trials. Fits for representative subjects are displayed in Supplementary Figures S2 and S3.

**Fig. 4:**
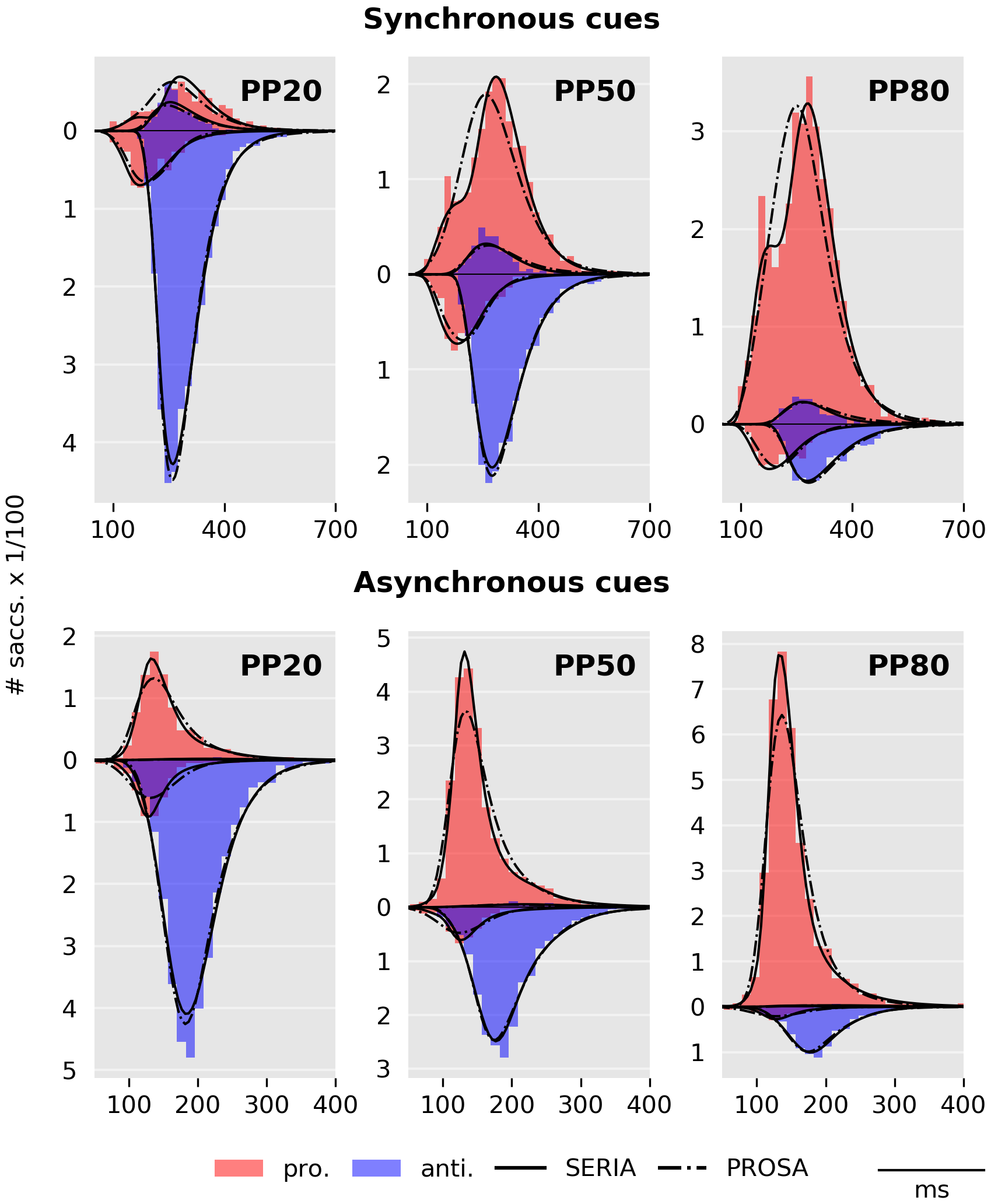
Histogram of RTs and model fits. For comparison, prosaccade trials are displayed in the positive half plane while antisaccades trials are displayed in the negative half plane. The histogram of prosaccade responses is displayed as red bars, whereas antisaccades are displayed in blue.

Replicating the findings in Aponte et al., 2017, the RT distribution of correct prosaccades in the SC condition was bimodal and could not be captured by the PROSA model, but was accounted for by the SERIA model. More importantly here, the RT distributions in the AC condition were also fitted better by the SERIA model. This was mostly clearly in correct prosaccades in the PP50 and PP80 condition (Fig. 4, bottom row, middle and right panels).

To further investigate the fits of the SERIA model, Fig. 5 displays the empirical and predicted cumulative density function (cdf). of the reciprocal RT of correct pro- and antisaccades. Cdf.’s are displayed in the probit scale (Noorani and Carpenter, 2016) but in contrast to Aponte et al., 2017 and Noorani and Carpenter, 2013, we did not normalize by the total number of saccades.

**Fig. 5:**
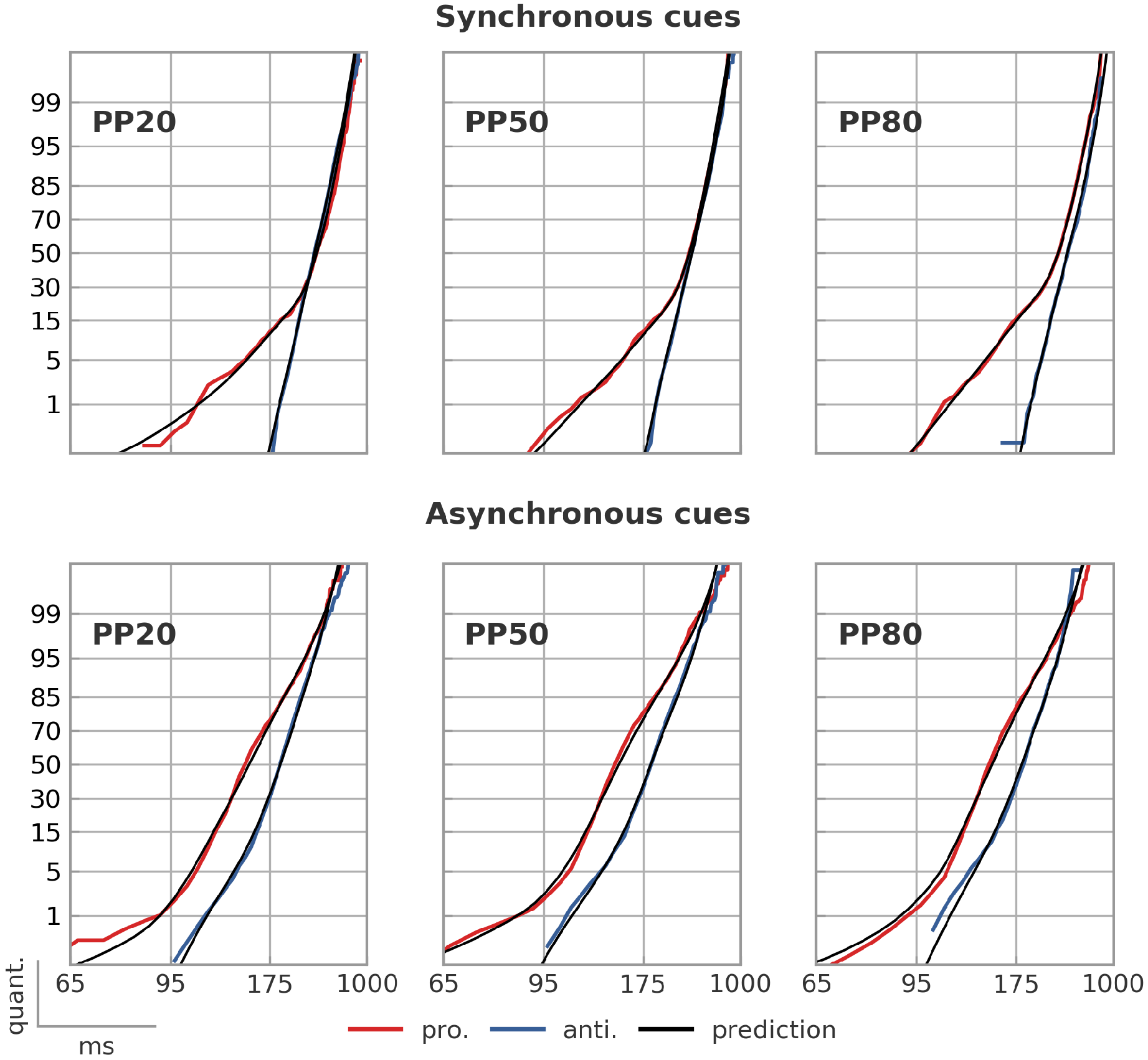
Empirical and predicted reciprobit of RTs in correct trials. In the SC condition, the SERIA model clearly captured the apparent bimodality of the RT distributions, although it did not completely account for the left tail of prosaccade RTs distribution in the PP20 and PP50 conditions. Please note the deflection in the prosaccade cdf, which demonstrates a bimodal distribution. In the AC condition, the SERIA model accounted for most of the relevant aspects of the RT distribution, e.g., left and right tails.

Reciprocal RTs in correct trials in the SC condition resembled the findings in Carpenter and Williams, 1995, and suggest that prosaccades are the results of two processes (Noorani and Carpenter, 2016). Moreover, the RT distribution of late prosaccades converges to the distribution of correct antisaccades. This provides further evidence for the conclusion (Aponte et al., 2017) that late prosaccades are the result of a slow accumulation process analogous to the one used to model antisaccades.

Importantly, SERIA also yielded accurate fits in the AC condition. Although the RT distribution of pro- and antisaccades deviated from the linear behavior observed in the SC condition, the model correctly predicted the empirical cdfs. Arguably, because late responses have latencies as low as 95ms, early and late prosaccades are disguised in a single unimodal distribution that does not follow the linear pattern observed in the SC condition. The PROSA model yielded less accurate fits (cf. Supplementary Figure S4).

### RT distribution of corrective antisaccades

Similarly as in Aponte et al., 2017, in order to predict the RT distribution of corrective antisaccades, the distribution of the hit time of the late antisaccade unit of each subject in each condition was weighted by the corresponding number of corrective antisaccades. The estimated distribution was time-shifted to optimize the predictive fit, i.e., we tried to predict the shape of the RT distribution, *not* its mean. Fig. 6 displays the predicted distributions in the SC (*time-shift=93ms*) and AC (*time-shift=63ms*) conditions. Visual inspection suggests that SERIA predicted correctly the shape of the distribution of corrective antisaccades.

**Fig. 6:**
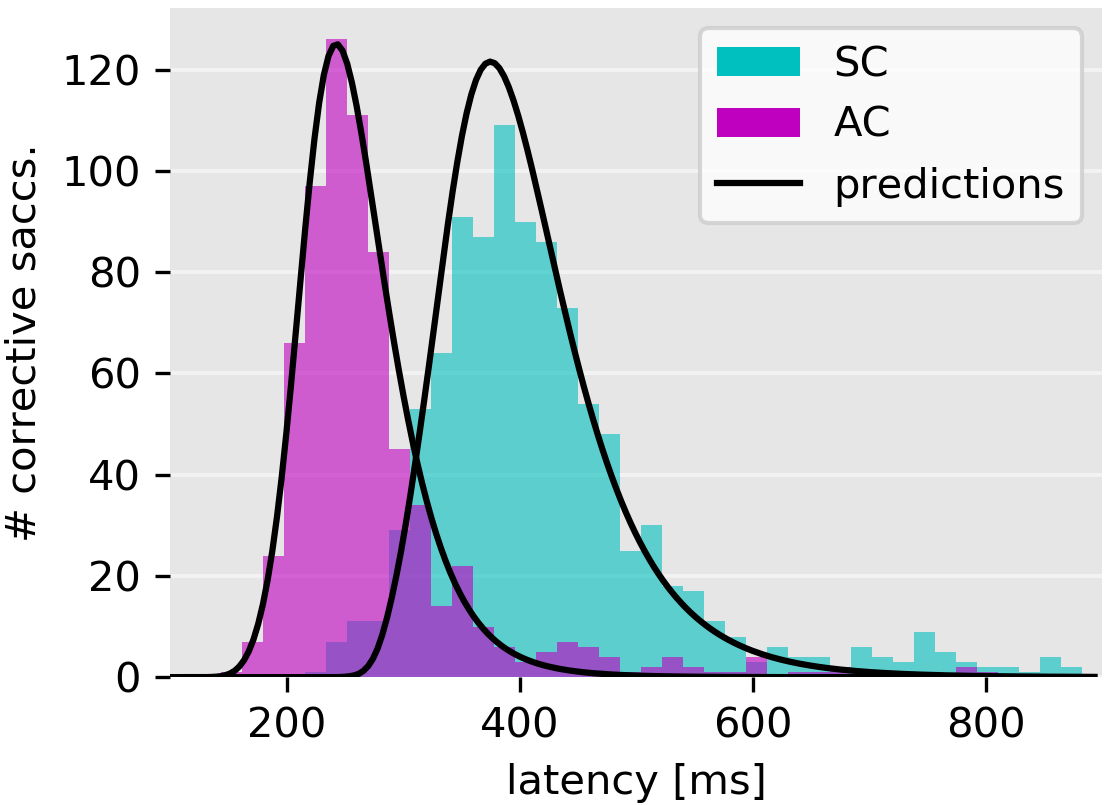
Histogram of corrective antisaccades and model predictions. Depicted are the distributions of the hit times of the antisaccade unit and the histogram of the corrective antisaccades’ RT. The location or time-shift of the predicted distributions was optimized using the data.

### Model parameters: Inhibition failures and late errors

We then turned our attention to inhibition and volitional or late errors. The latter are defined by the probability that the late prosaccade unit hits threshold before the antisaccade unit in an antisaccade trial, and the probability that the antisaccade unit hits threshold before the late prosaccade unit in a prosaccade trial. We also investigated the probability of an inhibition failure, i.e., the probability that the early unit hits threshold before all other units. In an antisaccade trial an inhibition failure corresponds to a reflexive error.

In the SC condition (Fig. 7A), our findings were similar to the results in Aponte et al., 2017. While the probability of a late error in a prosaccade trial was negatively correlated with PP (X^2^(2, *N* = 72) = 156.66, *p* < 10^−5^), the opposite behavior was observed in the probability of an inhibition failure in antisaccade trials (X^2^(2, *N* = 72) = 22.5, *p* < 10^−3^) and late errors in antisaccade trials (X^2^(2, *N* = 72) = 23.50, *p* < 10^−5^).

**Fig. 7:**
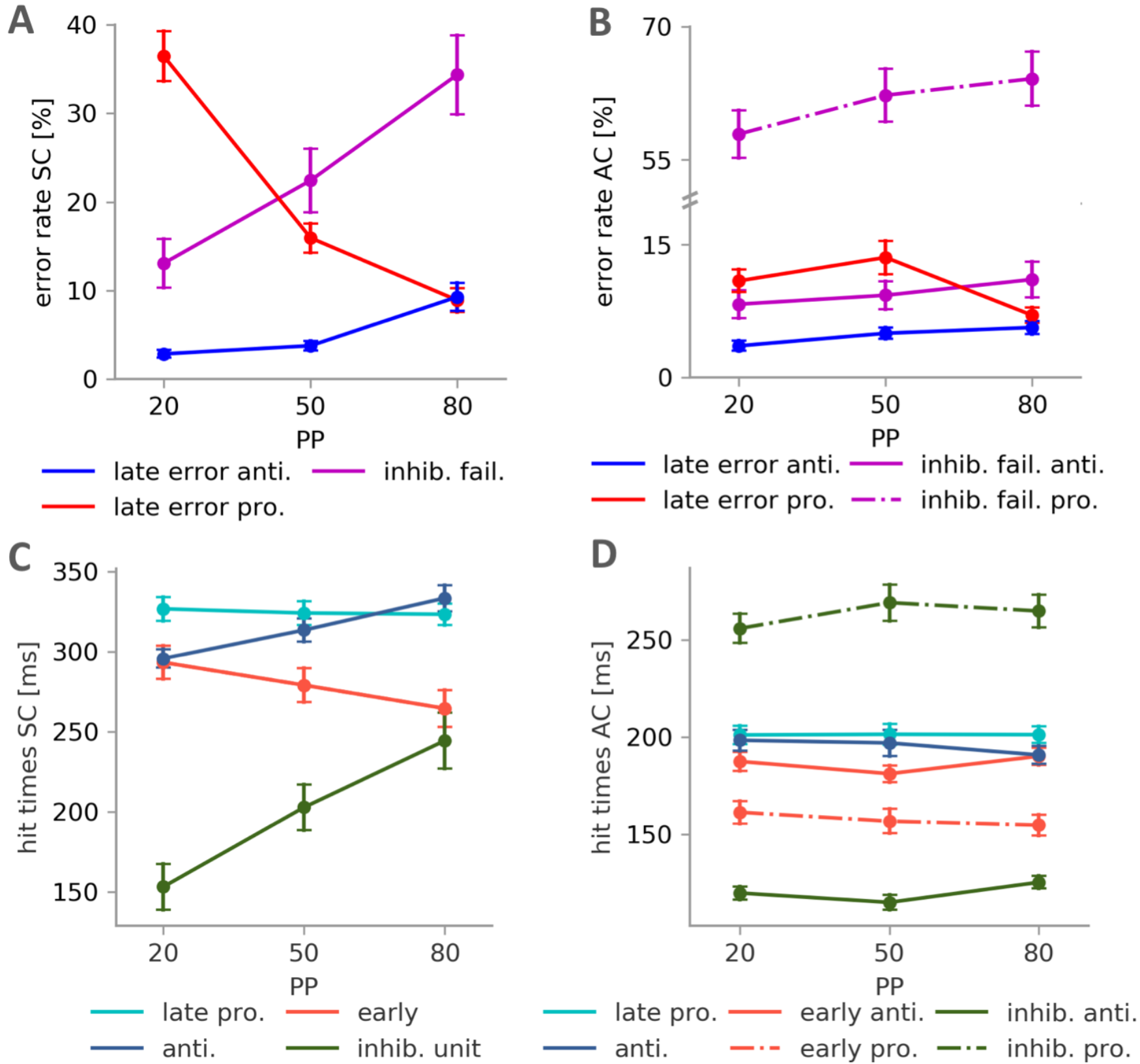
**A. Probability of late errors and inhibition failures in the SC condition.** Late errors occur when an early prosaccade is stopped by the inhibitory unit, but the incorrect late action is performed. Non-stopped early reactions are called inhibition failures. **B. Probability of late errors and inhibition failures in the AS condition. C. Expected hit time of the units in the SC condition.** Note that we report a single estimate for the early and inhibitory unit because in a constrained model both units are assumed to have the same behavior across trial types. **D. Expected hit time of the units in the AC condition.**

By contrast, in the AC condition it was necessary to consider the number of inhibition failures in pro- and antisaccade trials separately because model comparison favored models in which the early and inhibitory units behaved differently across trial types. We found (Fig. 7B), that the probability of an inhibition failure in prosaccade trials (mean 61%, std. 11) was much higher than in antisaccade trials (mean 9%, std. 8), indicating that most prosaccades were early, reflexive responses. When we considered the effect of PP in the AC condition, we found only a significant effect on the probability of a late error in antisaccade trials (X^2^(2, *N* = 72) = 6.31, *p* = 0.04).

In the AC condition, the percentage of late responses in prosaccade trials was estimated to be approximately 39% of all trials (see Fig. 7B and Table 3). In antisaccade trials, the percentage of inhibition failures was estimated to be 9% of the trials whereas the percentage of errors that could be attributed to the late decision process was 39%. In the SC condition, the number of antisaccade errors predicted by the model was approximately 2% higher than the empirical error rate. On average 21% of the errors were cataloged as late decision errors. To assess the posterior predictions of the model, we report the correlation coefficient between the empirical and predicted ER in Table 3.

**Table 3.** Empirical and predicted error, inhibition failures, and late errors. In order to evaluate the error rate estimates, we display the correlation coefficient between the predicted and observed error rates. Please note that inhibition failures in prosaccade trials correspond to correct early prosaccades. Errors in prosaccade trials can only be explained as late, volitional errors.

|  | Empirical and fitted error rates |  |  |  |  |  |
| --- | --- | --- | --- | --- | --- | --- |
|  | PP20 | PP50 | PP80 | PP20 | PP50 | PP80 |
|  | Antisaccade trials |  |  |  |  |  |
|  | SC |  |  | AC |  |  |
| Empirical error rate [%] | 14.06 | 22.79 | 38.22 | 11.56 | 13.81 | 16.83 |
| Predicted error rate [%] | 15.18 | 24.90 | 40.53 | 11.50 | 13.82 | 16.07 |
| Correlation coefficient | 0.99 | 0.97 | 0.98 | 0.99 | 0.94 | 0.99 |
| Inhib. failures [%] | 12.65 | 22.00 | 34.63 | 8.27 | 9.28 | 11.06 |
| $\frac{100 * \text{Late errors}}{\text{Late errors} + \text{inhib.fail.}}$ | 26.82 | 18.62 | 22.13 | 39.44 | 40.01 | 37.28 |
| Prosaccade trials |  |  |  |  |  |  |
| Empirical error rate [%] | 29.11 | 11.18 | 4.88 | 3.07 | 3.07 | 1.07 |
| Predicted error rate [%] | 30.91 | 11.70 | 5.21 | 3.13 | 3.14 | 1.12 |
| Correlation coefficient | 0.98 | 0.96 | 0.99 | 0.97 | 0.98 | 0.90 |
| Inhib. failures [%] | 13.08 | 22.42 | 34.34 | 57.86 | 62.23 | 64.10 |

We then investigated whether the percentage of inhibition failures in the SC condition was correlated with the percentage of inhibition failures in antisaccade trials in the AC condition. Results are displayed in Fig. 8. In each of the PP conditions, we found a significant correlation (*p* < 0.005) indicating that the tendency of individual subjects to respond with an early saccade was proportional across task designs.

**Fig. 8:**
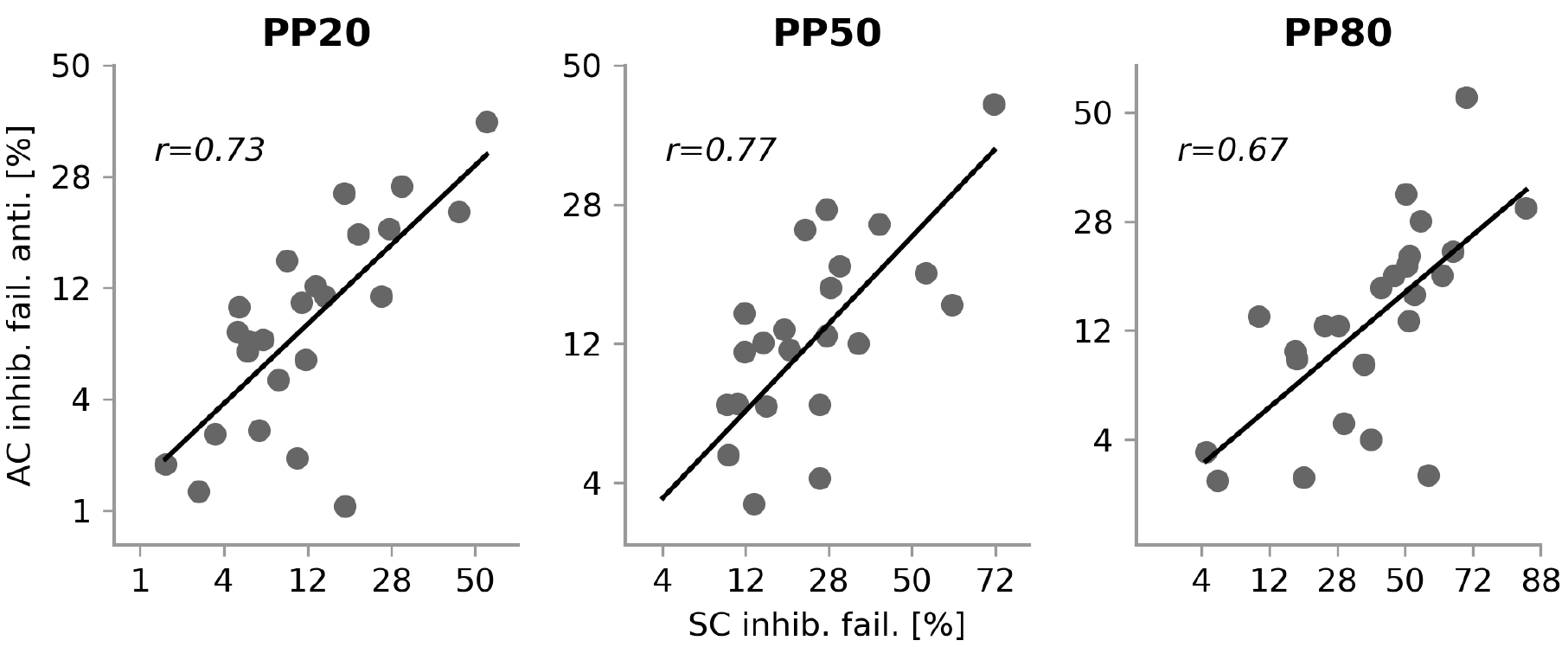
Correlation of inhibition failures in antisaccade trials. Values are displayed in the probit scale. There was a significant and very strong correlation between the percentage of inhibition failures across task designs and conditions.

### Model parameters: Hit times

To conclude, the effect of PP on the expected hit times of the units was investigated. In the SC condition (Fig. 7C), the early (*F*_2,46_ = 7.39, *p* = 0.001), as well as the antisaccade (*F*_2,46_ = 36.34, *p* < 10^−5^) and inhibitory units (*F*_2,46_ = 18.12, *p* < 10^−5^) were significantly affected by PP: High prosaccade trial probability led to slower inhibition, slower antisaccades, and faster early responses. However, we did not find a significant effect of PP on the hit times of the late prosaccade unit (*F*_2,46_ = 0.22, *p* = 0.79).

In the AC condition (Fig. 7D), most of the units had a much shorter hit time compared to the SC condition. Moreover, the fitted parameters suggested that most differences between pro- and antisaccade trials could be attributed to changes in the hit time of the inhibitory unit, which was over 100ms higher in prosaccade trials than in antisaccade trials. To further test this observation, we fitted a mixed Gamma SERIA model in which the early prosaccade unit (but not the inhibitory unit) was set to be equal across trial types. This is analogous to the restricted model originally proposed by (Noorani and Carpenter, 2013). This model obtained the highest evidence in the AC condition (Δ*LME* > 7 log units). Crucially, this restricted model was also better than one in which the early unit but not the inhibitory unit was allowed to change across trial types (*ΔLME* > 80). Thus, most variance in the probability of early prosaccades could be explained by changes in the inhibitory unit, which indicates that cuing the trial type in advance of the saccade direction cue mainly influenced the inhibition of early responses.

There was no significant effect of PP on the hit time of the late pro- and antisaccade units (late pro: *F*_2,46_ = 0.00, *p* = 0.99; anti: *F*_2,46_ = 2.08, *p* = 0.13). However, we found a significant effect of PP on the inhibitory unit regardless of the trial type (pro. trials: *F*_2,46_ = 3.23, *p* = 0.04; anti. trials: *F*_2,46_ = 14.11, *p* < 10^−3^). Finally, there was a significant effect of PP on the early unit in antisaccade trials (*F*_2,46_ = 8.62, *p* = 10^−3^), but not on prosaccade trials (*F*_2,46_ = 2.15, *p* = 0.12). Taken together, our results suggest that manipulating the trial type probability in AC task had only an effect on the early and inhibitory units, and this effect was weak in prosaccade trials.

## Discussion

There are four main findings in the present study. First, the SERIA model accounted for RT and ER better than the PROSA model in both the SC and AC conditions. This indicates that even in AC designs, the prosaccade RT distribution is best described by more than one decision process. Second, according to the model fits, a significant proportion of errors in antisaccade trials were late errors, irrespective of the CUE condition. Third, we found that in the AC condition, the main factor explaining the differences in ER and RT between pro- and antisaccade trials was the hit time of the inhibitory unit, and, with it, the probability of inhibiting an early response. Finally, we found that the effects of manipulating the probability of a trial type were almost completely abolished when subjects were cued about task demands in advance of the peripheral cue. Moreover, all effects of trial type probability were restricted to the early and inhibitory unit in the AC condition. We proceed to discuss these findings.

### The SERIA model accounts for antisaccade behavior regardless of CUE condition

Arguably, the main novelty of the SERIA model is the distinction between early responses, which are always directed toward a peripheral cue and can be inhibited by a stop process, and voluntary, late responses which can trigger both pro- and antisaccades. The units that trigger this type of saccades can generate rule guided behavior (e.g., an antisaccade), at the cost of higher RTs. Moreover, voluntary saccades are also subject to a race-to-threshold decision process, as shown in Aponte et al., 2017. This simple mechanism represents a unified explanation for late errors in antisaccade trials and errors in prosaccade trials.

In contrast, involuntary and voluntary saccades are often distinguished by the paradigm in which these are elicited (Walker et al., 2000) and not by the mechanism that generates them: On one hand, involuntary saccades are associated with paradigms in which a suddenly displayed stimulus elicits a saccade. On the other hand, voluntary saccades are associated with paradigms in which the target needs to be retrieved from memory or it depends on specific task instructions, such as in the antisaccade task.

Because the SERIA model allows both reflex like and ‘voluntary’ saccades towards a visual cue, the distinction between voluntary and involuntary saccades can be reformulated in terms of the processes that generates them. Accordingly, the antisaccade ‘cost’ (Hallett, 1978) might be also understood as a ‘voluntary’ saccade cost (ignoring remapping costs). This reconceptualization might explain the finding that under certain circumstances pro- and antisaccades exhibit the same latency (Liu et al., 2010, Weiler and Heath, 2014); if all early responses are inhibited, pro- and antisaccades could have the same latency.

Qualitatively, evidence for the SERIA model can be easily observed in the SC condition: correct prosaccades follow a bimodal distribution, and their late component resembles the distribution of correct antisaccades. Moreover, errors in prosaccades trials are relatively common in this version of the antisaccade task, and their latency is similar to the latency of correct antisaccades.

The main question that we addressed in this study is whether a similar mechanism could explain RT and ER distributions in an AC task design. Although correct prosaccade RTs do not show a bimodal distribution and errors in prosaccade trials are rare (<4%), model comparison and qualitative checks clearly indicate that prosaccade RT distributions in the AC condition can be better explained by a model that postulates early and voluntary prosaccades. Moreover, this model can predict the RT distribution of corrective antisaccades with surprising accuracy in both conditions.

Our data supports the idea that prosaccades do not appear to be bimodally distributed in the AC condition because voluntary prosaccades are fast enough to overlap with early prosaccades. This is obvious in Fig. 5 bottom row, in which the distribution of correct prosaccades deviates from the linear pattern usually observed in other conditions (see Fig. 5 top row and Noorani and Carpenter, 2016).

### Early and late errors in antisaccade trials

SERIA provides a formal account of errors in the antisaccade task which distinguishes it from two other prominent models. On one hand, the model in Noorani and Carpenter, 2013 does not incorporate a late decision process and thereby it explains all errors as inhibition failures. On the other hand, lateral inhibition models (Cutsuridis et al., 2007; 2014; Cutsuridis, 2015) explain errors as the result of connected accumulators that represent pro- and antisaccades, without the intervention of a third inhibitory unit. Accordingly, an error occurs when a voluntary action does not inhibit a reflex-like prosaccade. Along this line, Reuter and colleagues (Reuter and Kathmann, 2004) have argued that deficits in the ability to initiate an antisaccade contribute to the elevated ER observed in patients with schizophrenia.

The SERIA model is closer to the idea proposed by Fischer et al., 2000; Klein and Fischer,), who extended the distinction between ‘express’ and ‘normal latency’ saccades to antisaccade errors. Although conceptually similar, Klein and Fischer, 2005 used only a time threshold to distinguish between the two types of saccades. In this context, SERIA offers a model-based, statistically sound separation between early and late errors that goes beyond simple thresholding of RTs.

Hence, an important conclusion from our analysis is that late errors are a significant fraction of all errors regardless of task design. Concretely, in the present sample, approx. 39% of the errors in antisaccade trials in the AC condition were quantified as late errors, with large variability across subjects (Fig. 8). This is of significance, as the ability to separate between early and late errors might be of relevance in computational psychiatry and future patient studies (Fischer et al., 2000; Heinzle et al., 2016; Lo and Wang, 2016; Coe and Munoz, 2017).

### AC vs. SC designs

The most obvious difference between the AC and SC conditions was an overall reduction in RT and ER in the AC task. This observation replicates the findings in Weber, 1995 and a more recent study by Weiler and Heath, 2014.

There are two main explanations for these differences. First, in the SC condition the mapping between a cue and an action can only be started once the peripheral stimulus is presented. Thus, one would expect robust inhibition of reactive saccades, that affords processing of the peripheral cue (Weber, 1995). Second, in the AC condition subjects could anticipate the presentation of the peripheral cue, because the task cue was always displayed for 700ms. Despite this general reduction in RT, ERs were lower in the AC condition when compared to the SC condition.

Model comparison suggests differences in the type of anticipatory preparation in the two tasks: whereas in the SC condition, the early and inhibitory unit followed a similar hit time distribution across trial types, this was not the case in the AC condition. Furthermore, a model in which the prosaccade unit was fixed across trial types obtained the highest model evidence, indicating that most of the differences in the number of early responses could be accounted for by changes in inhibitory control.

One interpretation of our findings is that in the SC condition, the peripheral cue does not influence the inhibition of early responses, because it is integrated too late in the decision making process to strongly affect the early and inhibitory units. Nevertheless, contextual information about trial type probability can be exploited by the participants to drive inhibitory control. Contrastingly, in the AC condition early prosaccade inhibition is almost entirely determined by the trial type cue, and only weakly modulated by the probability of a trial type, as discussed below.

Importantly, the probability of antisaccade errors was correlated across both CUE conditions. Thus, relative ERs were consistent across the two tasks, suggesting that the same cognitive processes are involved in both conditions. In conclusion, SC designs are likely to provide more variability in terms of ER and RT, while probing the same cognitive processes involved in an AC paradigm.

### The effect of trial type probability

Our results replicate the finding that in the SC condition the probability of a trial type has a large impact on both ER and RT (Chiau et al., 2011; Aponte et al., 2017). Concretely, RTs of correct responses were negatively correlated with the corresponding trial type probability. These effects were strongly reduced in the AC condition, as reported before (Massen, 2004; Pierce et al., 2015; Pierce and McDowell, 2016a). Modeling indicated no significant effect of PP on late responses and a significant but relatively small effect on the early and inhibitory units.

### Summary

This study aimed to test whether and to what extent cue presentation order (task cue and spatial cue) influenced ER and RT in the antisaccade task. Overall, we found that the impact of trial type probability was strongly reduced in the AC condition compared to the SC condition. From a modeling perspective, our results demonstrate that the combination of an early and a late race between voluntary pro- and antisaccades better accounts for RT and ER in an AC design, as compared to models that incorporate only an early race. Furthermore, modeling revealed that early inhibitory processes are strongly influenced by trial type in the AC condition, but not in the SC condition. By contrast, trial type probability had a strong effect on early units in the SC condition, but not in the AC condition. SERIA also provided a good prediction of the shape of the distribution of corrective antisaccades in both tasks. Finally, our quantitative analysis supports the hypothesis that a non-negligible fraction of errors in the antisaccade task can be categorized as late errors.

### Software note

The models used here are available under the GPL licenses as part of the TAPAS toolbox (https://translationalneuromodeling.github.io/tapas/).

## Supplementary Material

### Supplementary Methods and Supplementary Figure S1

To infer the model parameters of all subjects we used the likelihood function defined by the SERIA and PROSA models and assumed a hierarchical prior, such that the parameters of all subjects in each CUE condition were estimated simultaneously. The graphical representation of the model is presented in Figure S1 following the conventions in Bishop, 2006.

**Figure S1:**
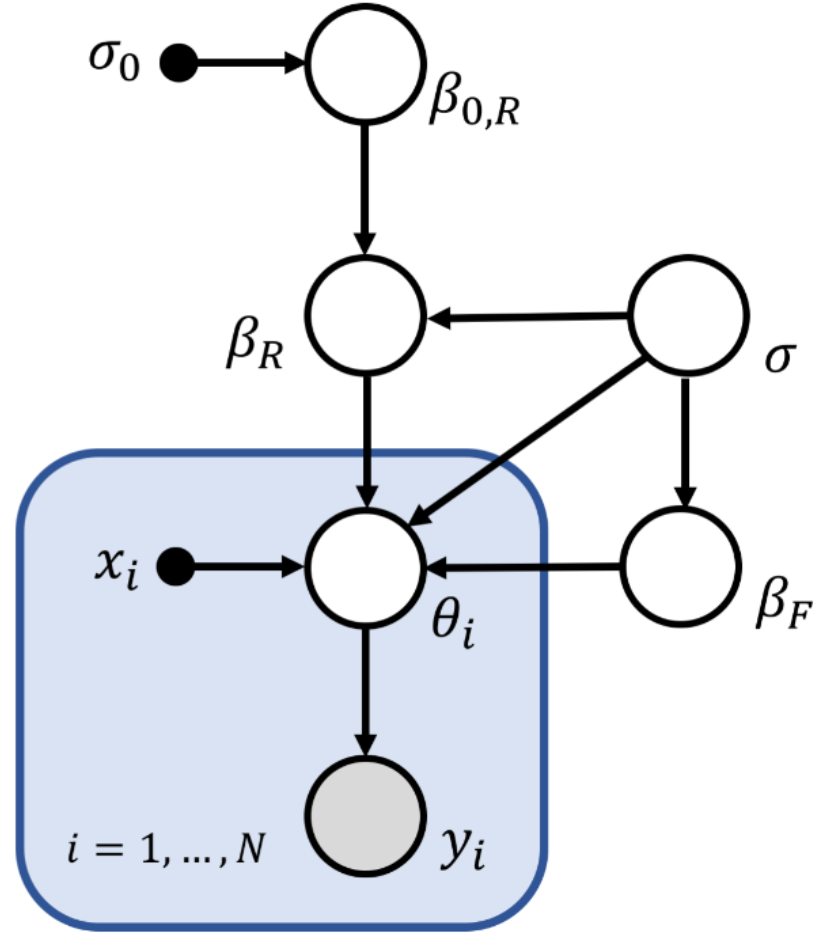
Graphical representation of the hierarchical model. The most important feature of the model is that the prior distribution of each set of parameters ***θ***_***i***_ is parametrically defined by a set of explanatory variables ***x***_***i***_ and coefficients ***β*** with variance ***σ***^**2**^. These coefficients are estimated from the population distribution. We partition parameters ***β*** into fixed (***β***_***F***_) and random effects (***β***_***R***_), such that the latter have a prior mean estimated again from the population distribution. For the present study, random effects represent subject specific intercepts, while their mean (or global intercept) is modeled by ***β***_**0, *R***_, whose prior distribution is assumed to be centered at zero with variance 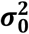.

To simplify notation, we assume that the data and parameters in each run are represented by a vector *y*_*i*_ and a single parameter *θ*_*i*_. The extension to a multivariate model is straight forward under the assumption that different parameters are conditionally independent. Although this assumption is likely to be inadequate, the main goal of the hierarchical extension of the model was to generate a data-driven prior mean and variance for the parameters, and not to account for the correlation between them. The likelihood of the model is given by the product of the likelihood of all runs:

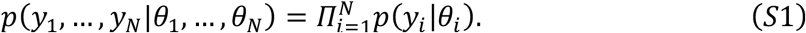

The prior distribution of parameters *θ*_*i*_ is given by

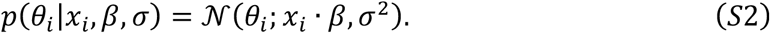

*X* = (*x*_1_,…, *x*_*N*_) is a design matrix of size *M* × *N* that codes *M* explanatory variables, such as SUBJECT, PP, etc., and *β* is a vector of dimension *M* × 1 that represents the effect of each explanatory variable. The prior distribution of *β* is given by

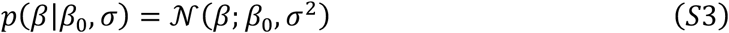

and the prior probability of *σ* is

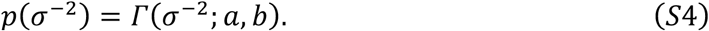

We distinguish between two types of *β* coefficients in analogy to the concepts of fixed and random effects. For fixed effects *β*_*F*_, we assumed that the coefficients have a fixed prior *β*_0, *F*_ = 0. For random effects *β*_*R*_, we assume that the prior mean *β*_0, *R*_ is a random variable such that

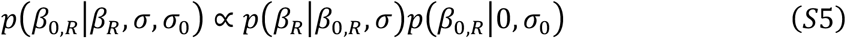

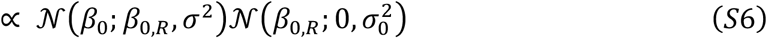

The conditional posterior of *β*_0, *R*_ is thereby

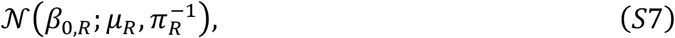

where

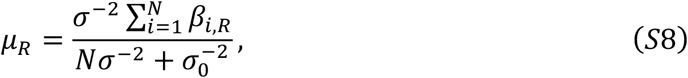

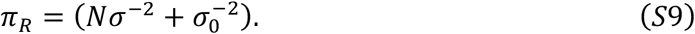

The rationale for including a random effect is to account for the idiosyncrasies of each subject individually while modeling a population wide intercept.

Since all the above equations are linear and rely on conjugate priors, it is possible to derive Gibbs steps to sample from the conditional posterior distributions of all parameters with the exception of *θ*_l_,…,_*N*_, which are sampled from a Gaussian kernel centered at the previous sample.

**Table S1:** Supplementary Table S1: Model comparison by accuracy. Expected log likelihood (accuracy) normalized by subtracting the lowest log likelihood (m2) from all estimates. The accuracy of a model is the expected log likelihood of the model. It is tightly related to the unpenalized R^2^, or total variance explained, of a model.

| **Model accuracy** |  |  |  |
| --- | --- | --- | --- |
| **Model** | **Parametric family** | **SC** | **AC** |
| **PROSA** |  |  |  |
| m<sub>1</sub> | T. normal | 224.6 | 338.9 |
| m<sub>2</sub> | T. normal | 0.0 | 0.0 |
| m<sub>3</sub> | Mixed Gamma | 607.1 | 376.3 |
| m<sub>4</sub> | Mixed Gamma | 499.0 | 72.4 |
| m<sub>5</sub> | Gamma | 613.2 | 347.8 |
| m<sub>6</sub> | Gamma | 531.2 | 126.1 |
| **DORA** |  |  |  |
| m<sub>7</sub> | T. normal | 1302.8 | 1011.0 |
| m<sub>8</sub> | T. normal | 1291.4 | 812.3 |
| m<sub>9</sub> | Mixed | 1351.4 | **998.7** |
| m<sub>10</sub> | Mixed | 1260.8 | 600.7 |
| m<sub>11</sub> | Gamma | **1390.9** | 893.0 |
| m<sub>12</sub> | Gamma | 1305.2 | 813.6 |

**Figure S2:**
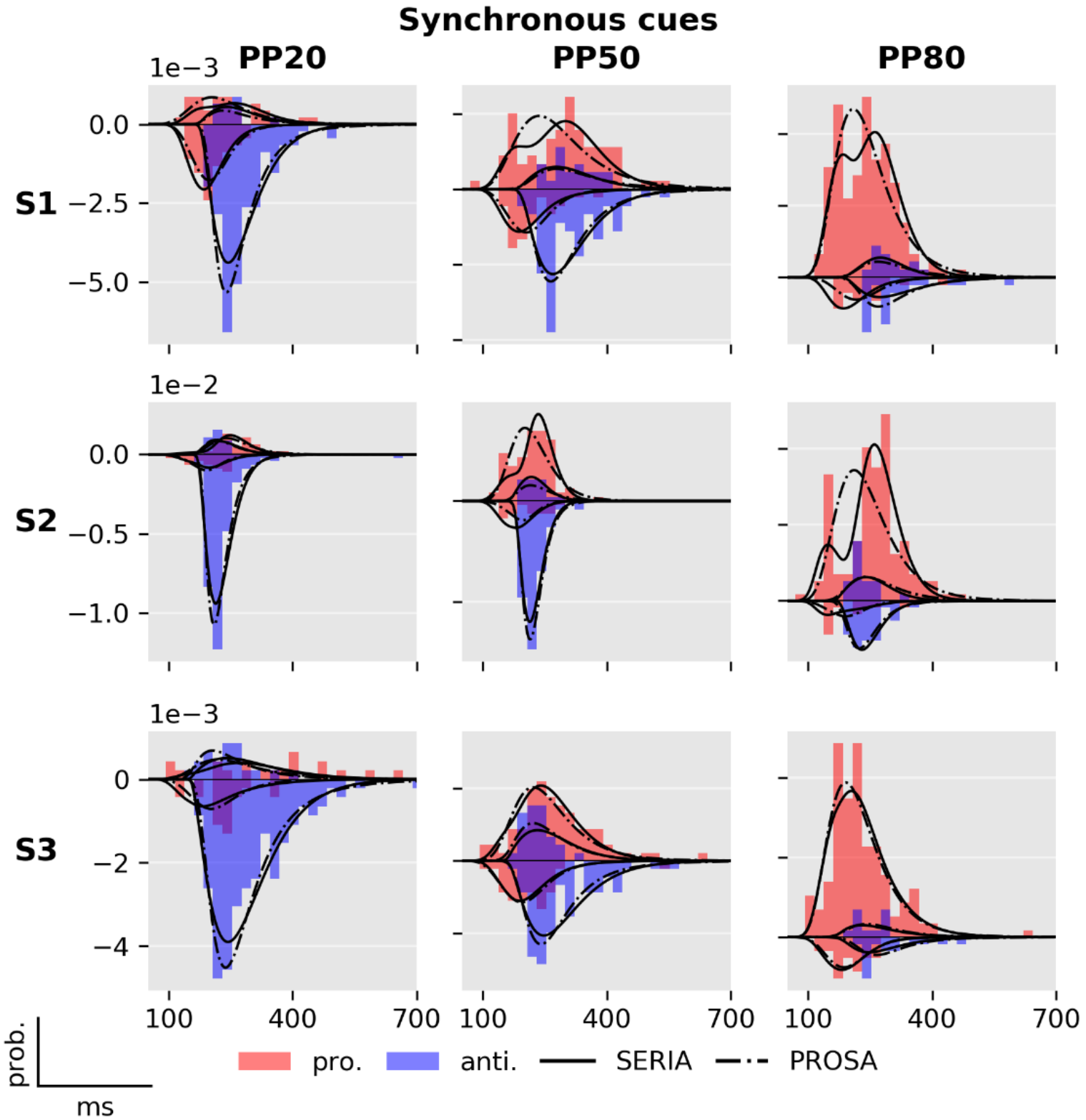
Supplementary Figure S2: Model fits for representative subjects - synchronous cue. RT distribution and comparison of the SERIA (m_10_) and PROSA (m_5_) models in the SC condition in three representative subjects, each one displayed in a different row. Prosaccade trials are displayed in the upper half plane, antisaccade trials in the bottom half. Note that the two models are the best models of their classes, respectively.

**Figure S3:**
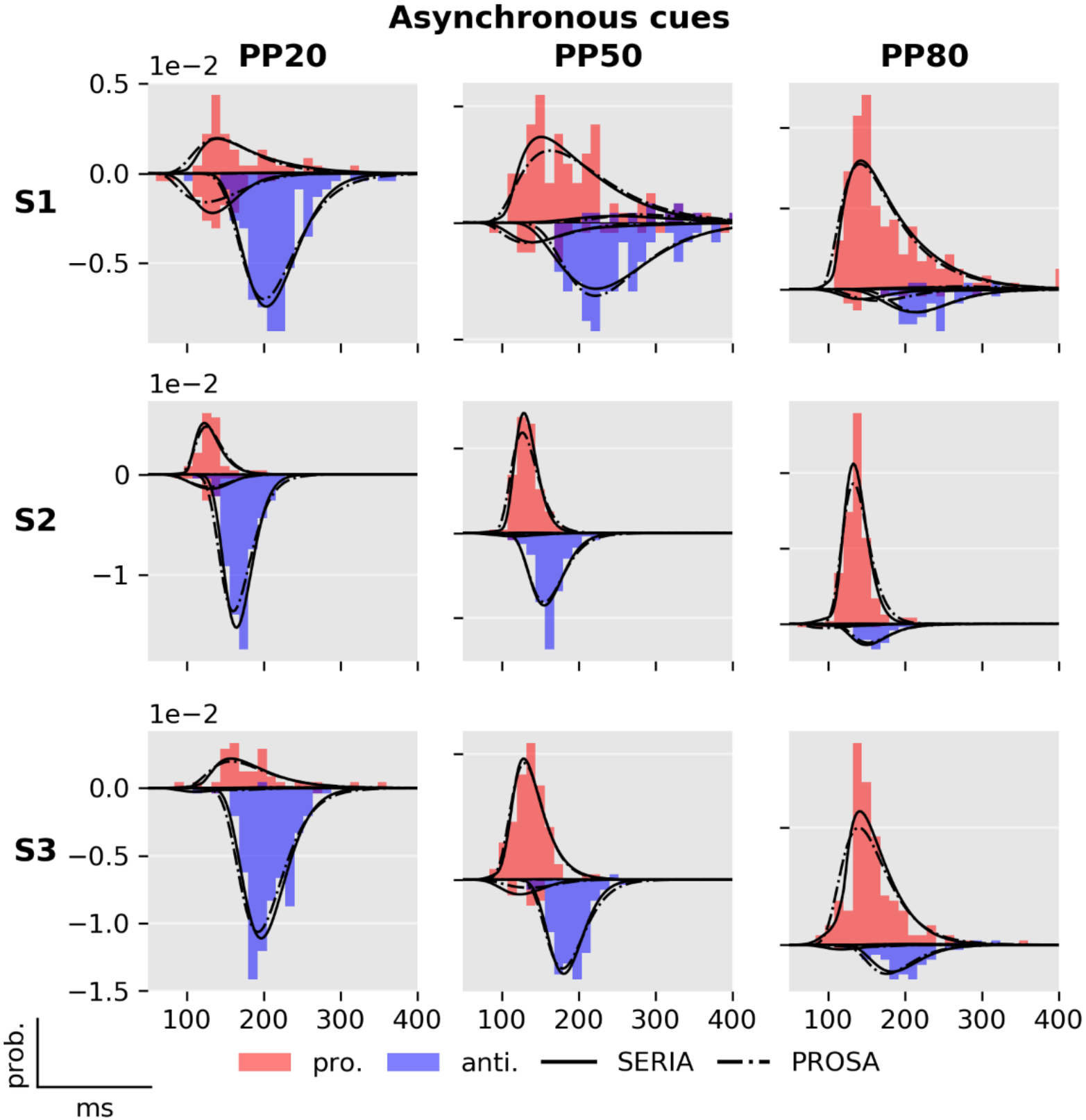
Supplementary Figure S3: Model fits for representative subjects - asynchronous cue. Comparison of the SERIA (m_9_) and PROSA (m_5_) models in the AC condition in three representative subjects, each one displayed in a different row. Note that the two models are the best in their respective families.

**Figure S4.:**
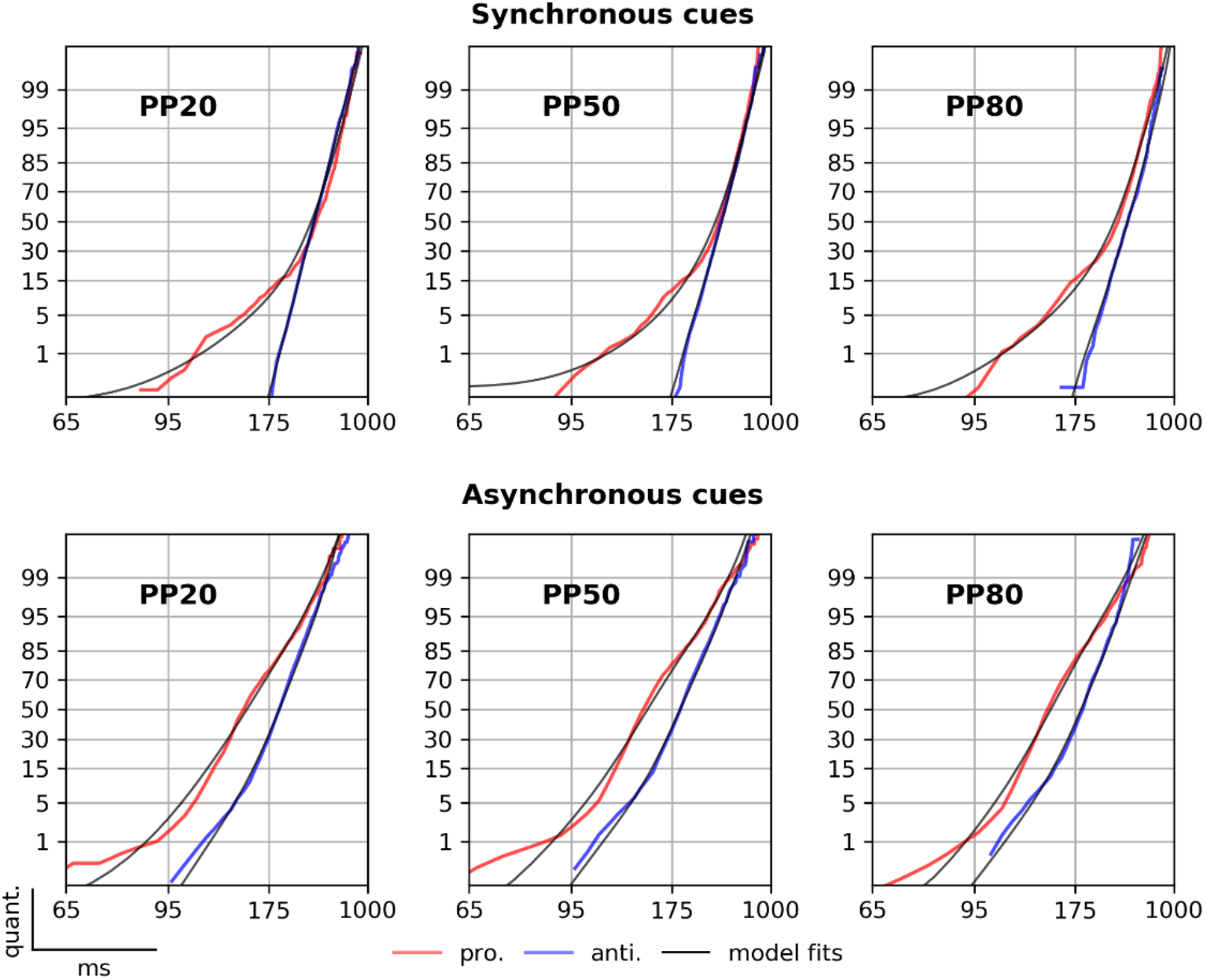
Supplementary Figure S4: Reciprobit plots of the PROSA model. Empirical and predicted reciprobit plots of the PROSA model. Same as in Figure 5, but using the PROSA model.

